# Amino acid sensing by the α-cell mitochondrial phosphoenolpyruvate cycle regulates intracellular Ca^2+^ levels independently of glucagon secretion

**DOI:** 10.1101/2025.05.26.656009

**Authors:** Erli Jin, Hannah R. Foster, Evgeniy Potapenko, Shih Ming Huang, Xinhang Dong, Jing W. Hughes, Matthew J. Merrins

## Abstract

Pancreatic islet α-cells are increasingly recognized as amino acid sensors for the organism. Building on our prior work in β-cells, we sought to determine whether the mitochondrial phosphoenolpyruvate (PEP) cycle is involved in α-cell amino acid sensing. Three different methods were used to probe the PEP cycle, including pyruvate kinase activators (TEPP-46), and mice with α-cell specific deletion of pyruvate kinase (PKM1/2-αKO) or mitochondrial PEP carboxykinase (PCK2-αKO). The mitochondrial fuels glutamine/leucine antagonized alanine/arginine-stimulated Ca^2+^ influx and glucagon secretion under hypoglycemic conditions. Both PKM1/2 and PCK2 were required for glutamine/leucine to close K_ATP_ channels and limit amino acid-stimulated membrane depolarization. The Ca^2+^ response to amino acids was suppressed by pyruvate kinase activation with TEPP-46, and enhanced by α-cell deletion of pyruvate kinase or PCK2 – all without changing glucagon secretion. Finally, using diazoxide/KCl to probe the pathways downstream of membrane depolarization, we identified an essential role of the PEP cycle in homeostatically restoring intracellular Ca^2+^ levels. In sum, the α-cell mitochondrial PEP cycle senses glutamine/leucine and inhibits K_ATP_ channels similarly to β-cells, while restricting amino acid-stimulated membrane depolarization and Ca^2+^ influx. However, defying expectations, none of the amino acids tested, including alanine/arginine, regulate glucagon secretion by modulating membrane depolarization or intracellular Ca^2+^.

Graphical abstract α-cells as amino acids sensors. Arginine, alanine, and glutamine potentiate glucagon secretion, while leucine has a suppressive effect (left). Independently of glucagon secretion, glutamine and leucine suppress alanine and arginine-stimulated Ca^2+^ influx via the phosphoenolpyruvate (PEP) cycle, which closes K_ATP_ channels and suppresses V_m_ depolarization (right).

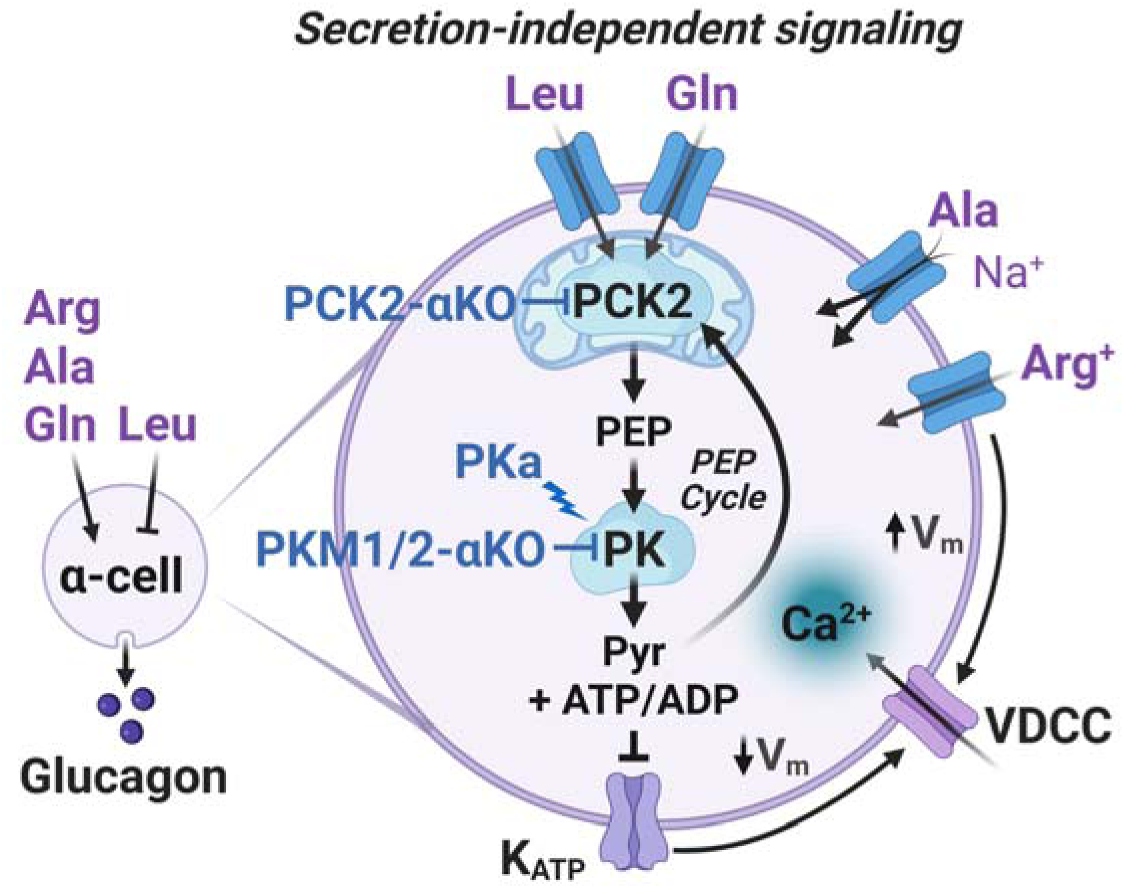

**Highlights:**

- Our studies identify a role for the α-cell PEP cycle in sensing amino acids under hypoglycemic conditions.
- Pyruvate kinase and PCK2 are required for glutamine/leucine to close α-cell K_ATP_ channel and limit membrane depolarization and Ca^2+^ influx.
- Glutamine/leucine oppose alanine/arginine-stimulated Ca^2+^ influx and glucagon secretion.
- All of the amino acids tested regulate glucagon secretion, but none do so by modulating membrane depolarization or intracellular Ca^2+^ levels.

## Introduction

Glucagon release from pancreatic islet α-cells is elevated under fasting and hypoglycemic conditions. Under these conditions, when inhibitory paracrine signals from neighboring islet β/δ-cells are reduced, α-cells respond strongly to non-glucose fuels including fatty acids and amino acids, which modulate Ca^2+^ and glucagon secretion (1–5). Glucagon secretion is dependent on voltage-dependent Ca^2+^ channels (VDCC) (6–8), which trigger downstream endoplasmic reticulum Ca^2+^ release (9). The activity of these channels is tightly regulated by membran potential (V_m_). Among the many ion channels that regulate α-cell V_m_, the activity of ATP-sensitive K^+^ (K_ATP_) channels is thought to be an important determinant of α-cell V_m_, Ca^2+^, and glucagon release (10–15). When glucose is elevated, glycolysis inhibits α-cell K_ATP_ channels and restricts Ca^2+^ entry (10,16–20). However, the metabolic pathways that control α-cell function in response to amino acid stimulation remain unknown.

How exactly do individual amino acids impact glucagon secretion? The mechanisms are diverse. Some amino acids, like glutamate and glycine, stimulate glucagon secretion through α-cell membrane receptors (21–23). Charged amino acids like arginine may stimulate Ca^2+^ influx by directly depolarizing V_m_ (24,25). Alanine, which is uncharged, may also regulate V_m_ due to co-transport with Na^+^ (26,27). In contrast, the branched chain amino acids are inhibitory to glucagon release (28–30). Leucine, for example, has been shown to suppress glucagon release by lowering cAMP, both via a direct effect on the α-cell and by stimulating inhibitory paracrine signaling via β/δ-cells (31). Beyond secretion, glutamine is required for amino acid stimulated α-cell proliferation (32), a mechanism that links glucagon release to amino acid catabolism in the liver-α-cell axis (3,33). Each of these findings indicate that amino acids are crucial to α-cell viability and function, but little is known about their intracellular metabolism.

In this paper, we focused on the mitochondrial fuels glutamine and leucine, and their control of α-cell V_m_ and Ca^2+^ by the glucagonotropic amino acids alanine and arginine. Due to the paucity of α-cell data, our working model was based on prior work in β-cells (18,34,35). At low glucose, glutamine and leucine evoke insulin secretion by stimulating mitochondrial phosphoenolpyruvate (PEP) production by the PCK2 enzyme. Mitochondria PEP is then used by plasma-membrane associated pyruvate kinase (PK), which raises the ATP/ADP ratio to close K_ATP_ channels, depolarize V_m_, and open VDCC (18,34–36). In the PEP cycle, pyruvate generated by the PK reaction is sequentially metabolized by mitochondrial pyruvate carboxylase and PCK2, and returned to cytosol as PEP to augment flux through PK. Similar to β-cells, α-cell K_ATP_ channels are locally controlled by plasma membrane-associated enzymes including PK (18). However, these experiments were performed using excised patch-clamp, leaving it unclear whether PK and the PEP cycle are active in α-cells and if they regulate V_m_, Ca^2+^, or glucagon secretion.

## Research Design and methods

### Mice

Wild-type C57BL/6J mice were obtained from The Jackson Laboratory. *GcgCre^ERT^* mice (Jax 030346) (46) were crossed with *GCaMP6s* mice (Jax 028866), a Cre-dependent Ca^2+^ indicator strain. *Pkm^f/f^* mice were generated using CRISPR and sequence-verified by the University of Wisconsin-Madison Genome Editing and Animal Model Core, as done previously to generate *Pck2^f/f^*mice (35). *Pkm^f/f^* and *Pck2^f/f^*mice were crossed with *GcgCre^ERT^:GCaMP6s* mice to generate *GcgCre^ERT^:GCaMP6s:Pkm^f/f^* (PKM1/2-αKO) and *GcgCre^ERT^:GCaMP6s:Pck2^f/f^* (PCK2-αKO) and their littermate *GcgCre^ERT^:GCaMP6s* controls. As in our prior study, *GcgCre^ERT^* was present in the dams to facilitate *loxP* excision in the study animals without the need for tamoxifen (31). Mice were used between 11-24 weeks of age. Although 4-hydroxytamoxifen (100 nM) was only strictly necessary for adenoviral expression of Cre-dependent JEDI-2P biosensors, it was included in the culture media in all islet studies for consistency, after verifying that hormone secretion was unaffected (Supplemental Fig. 6). All procedures involving animals were approved by the Institutional Animal Care and Use Committees of the William S. Middleton Memorial Veterans Hospital and the University of Wisconsin-Madison and followed the National Institutes of Health Guide for the Care and Use of Laboratory Animals.

### In vivo assays

For meal tolerance tests, mice were fasted for 5 hours prior to oral gavage of liquid Ensure (10 mL/kg). For insulin tolerance tests, animals were fasted for 5 hours, followed by IP-injection of 0.75 mg/kg Humilin-R (Eli Lilly). Blood glucose was measured from the tail using a glucometer (Contour). Glucagon was assayed from blood plasma using a mouse glucagon ELISA kit (Crystal Chem 81518) assayed with a TECAN Spark plate reader.

### Islet Perifusion Assays

Islets from 5-6 mice of each genotype were pooled and then divided in equal numbers (80-100 islets/chamber) and placed into a 12-channel perifusion system (BioRep Peri-5 or homemade perifusion system) containing KRPH buffer (in mM: 140 NaCl, 4.7 KCl, 2.5 CaCl_2_, 1 NaH_2_PO_4_, 1 MgSO_4_, 2 NaHCO_3_, 5 HEPES, 2 glucose, 0.1% fatty acid-free BSA, pH 7.4) using 100 μL Bio-Gel P-4 Media (Bio-Rad #1504124) to stabilize the islets. Islets were equilibrated at 2 mM glucose for 48 min prior to perfusion with AA at 37°C. Glucagon secretion was assayed using Promega Lumit Glucagon Immunoassay (W8020) and measured using a TECAN Spark plate reader. The Quant-IT PicoGreen dsDNA Assay Kit (Invitrogen P7589) was used to determine DNA content after lysis in 0.1% Triton X-100.

### Immunofluorescence staining and quantification

Isolated islets were fixed in 4% paraformaldehyde, permeabilized in CAS-Block (Invitrogen) containing 0.3% Triton X-100 for 1 hour at room temperature, and subsequently incubated overnight at 4°C with primary antibodies diluted in CAS-Block/Triton: rabbit-anti-PKM1/2 (CST3190T, 1:200); rabbit-anti-PCK2 (CST6924S, 1:200); mouse-anti-glucagon (ab10988, 1:800). Following five 10-minute washes in PBST (PBS with 0.1% Triton X-100), islets were incubated with secondary antibodies (Abberior STAR Orange anti-mouse and STAR Red anti-rabbit, 1:200) for 4 hours at room temperature, followed by five 10-minute PBST washes. Stained islets were mounted on glass slides using ProLong Gold Antifade (Invitrogen) and imaged on a Nikon AXR NSPARC confocal microscope. Z-stacks were acquired at 1 µm step size, covering 10-30 planes per islet. For quantification, glucagon signal was used to define α-cell vs. non-α-cell regions of interest (ROI). Whole-cell PKM1/2 and PCK2 staining intensities were measured within randomly selected ROIs across z-planes and islets. Statistical comparisons were performed using ordinary one-way ANOVA.

### FACS sorting and qPCR

Islets isolated from 6mice/group were pooled and dispersed with trypsin (TrypLE, Thermo fisher). Cells were sorted using FACSAriaII at University of Wisconsin Carbone Cancer Center Flow Lab. α and non α-cells were sorted based on GCaMP6s signal. RNA was isolated using QIAGEN RNeasy mini kit according to the manufacturer’s instructions. cDNA was prepared using monkey Moloney virus reverse transcriptase (Life Technologies) then analyzed by quantitative PCR (qPCR) using Power SYBR Master Mix (Life Technologies)(Forward primer: TGGCCGTGCAATCCAGAAAA; Reverse primer: TTGGTGATGCCCAAAATCAGCAT). All transcripts were normalized to β-actin. Fold changes were calculated relative to vehicle only controls.

### Electrophysiology

Mouse islets were dispersed with Accutase (Sigma-Aldrich A6964100ML) and plated on sterilized uncoated glass shards and cultured for up to 3 days. Recording electrodes made of microfilament borosilicate glass (Harvard Apparatus) were used to pull ∼3 MΩ patch pipettes using a Flaming/Brown micropipette puller (P-1000, Sutter Instruments), and polished (Narishige MF-830) to a final tip resistance of 4-5 MΩ. Cell-attached recording started after formation of a stable gigaseal (>3 GΩ). Recordings were made by using HEKA Instruments EPC10 patch-clamp amplifier at room temperature. The pipette solution contained (in mM): 10 sucrose, 130 KCl, 2 CaCl_2_, 0.9 MgCl_2,_ 10 EGTA, 20 HEPES, pH 7.2, adjusted with KOH. The extracellular bath solution contained (in mM): 140 NaCl, 5 KCl, 1.2 MgCl_2_, 2.5 CaCl_2_, 0.5 glucose, 10 HEPES, pH 7.4, adjusted with NaOH. Holding membrane potential was set at −50 mV during recording. Data was filtered online at 1 kHz with a Bessel filter and analyzed offline using Clampfit 10.7 software (Molecular Devices). During each condition, recordings are held for several minutes for the activity to stabilize, such that no significant further changes in channel conductance were observed. A 120-s window at the end of each condition was used to analyze K_ATP_ activity in terms of power, frequency, and duration and open time as previously described (18), all of which were normalized to control conditions. Membrane potential measurement was performed as in(47). Briefly, Intact islets were perfused with a standard external solution containing (135 mM NaCl, 4.8 mM KCl, 5 mM CaCl_2_, 1.2 mM MgCl_2_, 20 mM HEPES, 10 mM glucose; pH 7.35) with 2mM glucose and amino acids as indicated. Pipette tips were filled with an internal solution (28.4 mM K_2_SO_4_, 63.7 mM KCl, 11.8 mM NaCl, 1 mM MgCl_2_, 20.8 mM HEPES, 0.5 mM EGTA; 40 mM sucrose; pH 7.2) containing 0.36 mg/mL amphotericin B. Islet α and β-cells were identified by their characteristic I-V curves (Supplemental Fig. 3)

### Islet Ca^2+^ and V_m_ imaging

Reagents were obtained from Sigma-Aldrich unless indicated otherwise. For imaging of α-cell Ca^2+^, islets isolated from *GcgCre^ERT^:GCaMP6s* mice were cultured in the presence of 100 nM 4-hydroxytamoxifen and imaged 3 days post isolation. Islets with different genotypes were barcoded 391 by preincubation with 2 µM DiR (Thermo Fisher Scientific D12731) in 2ml islet media for 15 min at 37 °C. Islet barcode was then imaged using Cy7 (for DiR) from Chroma.For imaging of α-cell membrane potential, islets from *GcgCre^ERT^* mice were incubated with adenovirus expressing Cre-dependent JEDI-2P biosensors (Ad-CMV-FlexOn-JEDI-2P, VectorBuilder) for 2 hours, transferred to fresh media containing 100 nM 4-hydroxytamoxifen, and imaged 3 days post isolation. Islets were imaged on an RC-41LP chamber (Warner Instruments) mounted on a Nikon Ti2 microscope equipped with a 10×/0.5 NA SuperFluor objective (Nikon). The imaging solution contained (in mM): 137 NaCl, 5.6 KCl, 1.2 MgCl_2_, 0.5 NaH_2_PO_4_, 4.2 NaHCO_3_, 10 HEPES, and 2.6 CaCl_2_ (pH 7.4). Metabolites, mouse GIP (Pheonix Pharmaceuticals), and pyruvate kinase activator (TEPP-46, Calbiochem) were added as indicated. Islets were imaged at a constant flow rate of 250 µL/min using a feedback-controlled flow cell (Fluigent) and the temperature was maintained at 37°C with chamber and solution heaters (Warner). Filters for GCAMP6s and JEDI-2P imaging were as follows: exciter, ET500/20x; dichroic, ET-Fura2-GFP; emitter, ET535/30m (Chroma). Images were captured with a Hamamatsu ORCA-Flash4.0 CMOS camera every 6 s. A single region of interest was used to quantify the average response of each islet using NIS-Elements software (Nikon Instruments). Ca^2+^ and membrane potential were reported as the fluorescence intensity normalized to the initial condition (F/F_o_). Single α-cell quantification was performed with the Imaris analysis software (Oxford Instruments).

## Results

### Glutamine and leucine suppress α-cell Ca^2+^ influx stimulated by alanine and arginine

Mixed amino acids, provided at their concentrations measured in mouse portal blood (QLRA; 0.6 mM glutamine, 0.5 mM leucine, 0.2 mM arginine, 2 mM alanine) (26), stimulated biphasic glucagon secretion from mouse islets under low glucose conditions (Fig. 1A). To determine how amino acids regulate α-cell Ca^2+^, we imaged islets isolated from mice expressing GCaMP6s biosensors in α-cells (*GcgCre^ERT^:GCaMP6s*). Although single α-cells exhibited heterogeneity in their Ca^2+^ response, the aggregate islet response to mixed amino acids was a sustained rise in Ca^2+^ (Fig. 1B). Opening K_ATP_ channels with 200 μM diazoxide (Fig. 1C) or blocking L-type VDCCs with 1 μM isradipine (Fig. 1D) suppressed the Ca^2+^ rise stimulated by amino acids, confirming the requirement for membrane depolarization and L-type VDCCs (6,9,15).

**Figure 1.**
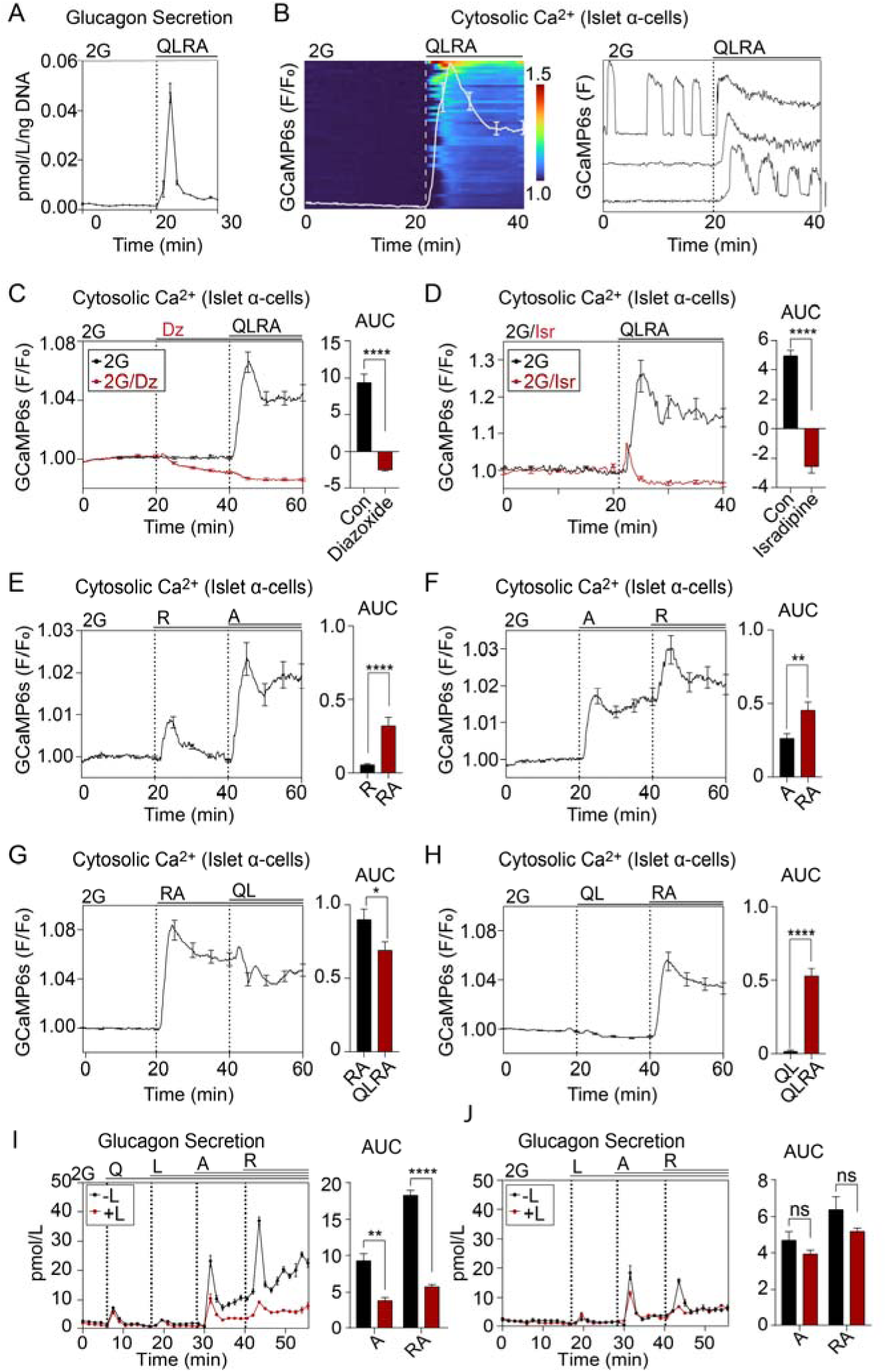
Glutamine/leucine suppress α-cell Ca^2+^ influx stimulated by alanine/arginine. *A*: Perifusion assay was used to measure glucagon secretion from mouse islets in the presence of 2 mM glucose (2G) and amino acids (QLRA, in mM: 0.6 mM glutamine, 0.5 mM leucine, 0.2 mM arginine, 2 mM and alanine). *B*: Heatmap (left) and example traces (right) of Ca^2+^ responses in intact mouse islets expressing GCaMP6s biosensors. Average Ca^2+^ is shown in the white trace (left panel). *C-D* Amino acid stimulated Ca^2+^ influx is blocked by 200µM diazoxide (K_ATP_ channel opener) (*C*) and 10 µM isradipine (L-type VDCC blocker) (*D*). *E-J*: α-cell Ca^2+^ (*E-H*) and glucagon secretion (*I-J*) stimulated by QLRA alone or in combination. Imaging data (*B*-*H*) represent averaged normalized GCaMP6s intensity from *n* = 40-56 islets from 3 mice imaged per protocol. Perifusion data (*A*, *I*, and *J*) represent the average of *n* = 3 chambers from *n* = 3 mice per condition with 80–100 mouse islets per chamber. Data are shown as mean ± SEM with AUC for each condition compared with control by Student’s *t* test. **\****P* < 0.05, **\*\****P* < 0.01, **\*\*\*\****P* < 0.0001.

We applied various combinations of amino acids at low glucose to identify their individual effects on α-cell Ca^2+^. Arginine induced only a transient Ca^2+^ rise, while alanine provoked a sustained increase (Fig. 1E,F). The effects of arginine and alanine were additive. On their own, the combination of glutamine and leucine only slightly reduced average Ca^2+^, however, they impaired the intracellular Ca^2+^ response to alanine and arginine by 42% (cf. Figs. 1G and H). In a perifusion assay, glutamine and leucine suppressed glucagon secretion stimulated by arginine and alanine (Fig. 1I). Glucagon release fell in response to glutamine withdrawal, and the glucagonostatic effect of leucine was reduced (Fig. 1J). Taken together, alanine and arginine stimulate Ca^2+^ influx and glucagon secretion, while the combination of glutamine and leucine has a suppressive effect. Since amino acids were applied to the entire islet in theses assays, the inhibitory effects of paracrine signaling cannot be completely ignored (*e.g.*, glutamine/leucine stimulate insulin release, Supplemental Fig. 1). We therefore sought additional approaches to isolate the effects of amino acids on α-cell metabolism.

### Mice lacking **α**-cell PKM1/2 or PCK2 respond normally to changing glucose levels

To test the function of the α-cell PEP cycle, we generated mice with α-cell specific knockout of PKM1/2 or PCK2 by crossing *GcgCre^ERT^:GCaMP6s* mice with *Pkm^f/f^*mice (PKM1/2-αKO, *GcgCre^ERT^:GCaMP6s:Pkm^f/f^)* or *Pck2^f/f^* mice (PCK2-αKO, *GcgCre^ERT^:GCaMP6s:Pck2^f/f^*). All studies were performed with littermate *GcgCre^ERT^:GCaMP6s* controls. α-Cell specific knockdown of PKM1/2 and PCK2 was evident by immunostaining of isolated islets (Fig. 2A,B), however high background fluorescence made absolute quantification difficult. After pooling FACS-sorted GCaMP6s^+^ α-cells from 6 mice per genotype, we confirmed by Western blot that PKM1 and PKM2 were virtually absent in α-cells from PKM1/2-αKO islets (Fig. 2C). Although PCK2 was undetectable by Western blot, we confirmed the loss of *Pck2* mRNA using QPCR of FACS-sorted α-cells; *Pck2* mRNA was reduced by 88% in PCK2-αKO α-cells relative to controls.

**Figure 2.**
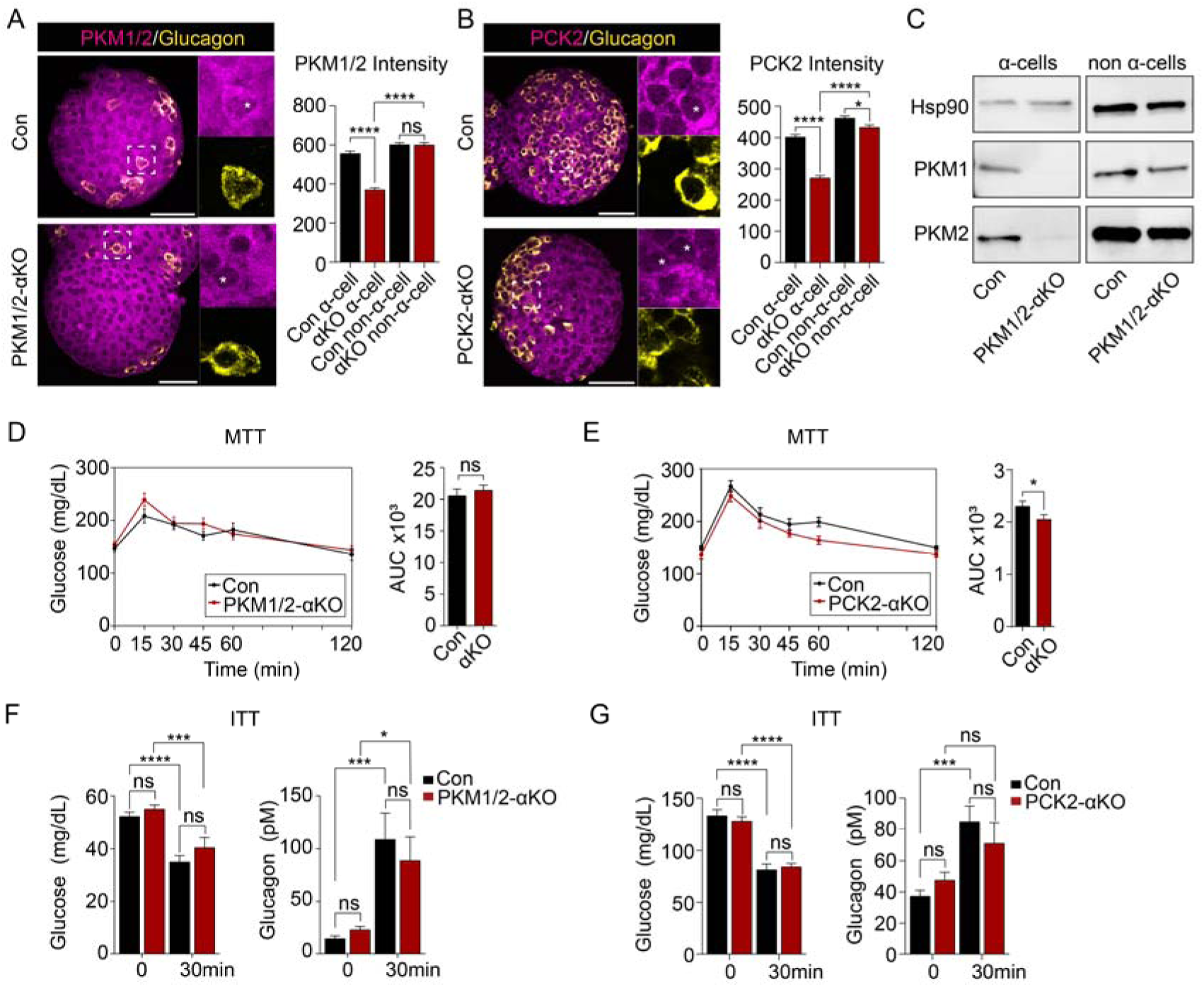
Mice lacking α-cell PKM1/2 or PCK2 respond normally to changing glucose levels. *A-B*: Representative images of stained islets showing targeted deletion of PCK2 in Glucagon^+^ α-cells. Insets highlight α-cells (asterisks) with intact (control) or reduced (PCK2-αKO) enzyme expression. Scale bar, 50 µm. Quantification represents *n* = 100 cells from 10 islets per genotype. C: Western blot showing PKM1/2 knockdown in FACS sorted α-cells (GCaMP6s^+^) and non-α-cells (GCaMP6s^-^) from PKM1/2-αKO and controls (pooled samples from 6 mice per group). D-E: Meal tolerance test (MTT) with PKM1/2 and PCK2 α-KO. *n* = 10 mice per group. *F-G*: Plasma glucose and glucagon levels in PKM1/2-αKO, PCK2α-KO, and littermate controls in response to an insulin tolerance test (ITT). Data are shown as mean ± SEM with AUC for each condition compared with control by Student’s *t* test (*C-D*) or one-way ANOVA (*B*, *E-F*). \**P* < 0.05, \*\*\**P* < 0.001, **\*\*\*\***P < 0.0001, ns, not significant.

To investigate glycemic control in the PKM1/2-αKO and PCK2-αKO mice, we performed meal tolerance tests (MTT) by gavage of protein shake after a 5 hour fast. No differences in blood glucose clearance were observed in the PCK2-αKO mice relative to controls (Fig. 2D). A small improvement in glucose tolerance was observed in the PCK2-αKO mice relative to controls, however this effect appears to be driven by elevated glucose in the control mice at a single timepoint (Fig. 2E). To test the function of PKM1/2-αKO and PCK2-αKO mice under low glucose conditions, fasted mice were subjected to an insulin tolerance test. No difference in fasting glucagon was observed in either strain, and insulin lowered blood glucose and stimulated glucagon secretion to a similar extent in PKM1/2-αKO and PCK2-αKO mice as controls (Fig. 2F,G). Thus, both strains of knockout mice responded normally to raising or lowering blood glucose.

### The **α**-cell PEP cycle closes K_ATP_ channels and restricts V_m_ depolarization in response to amino acid stimulation

Previous studies in β-cells show that, at low glucose, glutamine and leucine stimulate insulin secretion by fueling the mitochondrial PEP cycle to close K_ATP_ channels, depolarize V_m_, and open VDCCs (35). Given that α-cell K_ATP_ channels are also regulated by PK (18), we tested whether the PEP cycle impacts electrical activity similarly in α-cells. K_ATP_ channel activity was directly measured by patch-clamp using the fluorescent GCaMP6s reporter to positively identify α-cells. K_ATP_ channel activity was increased by glutamine and reduced by leucine in the control group, effects that were blunted or blocked by α-cell deletion of PKM1/2 and PCK2 (Figs. 3A and 3B). Notably, leucine retained some ability to close K_ATP_ channels in the PKM1/2-αKO, while failing to close K_ATP_ channels in the PCK2-αKO cells. These data suggest that the remaining α-cell PKL isoform can partially compensate for the loss of PKM1/PKM2, while cutting off the mitochondrial PEP supply for all three PK isoforms (i.e., by deleting PCK2) is more severe.

**Figure 3.**
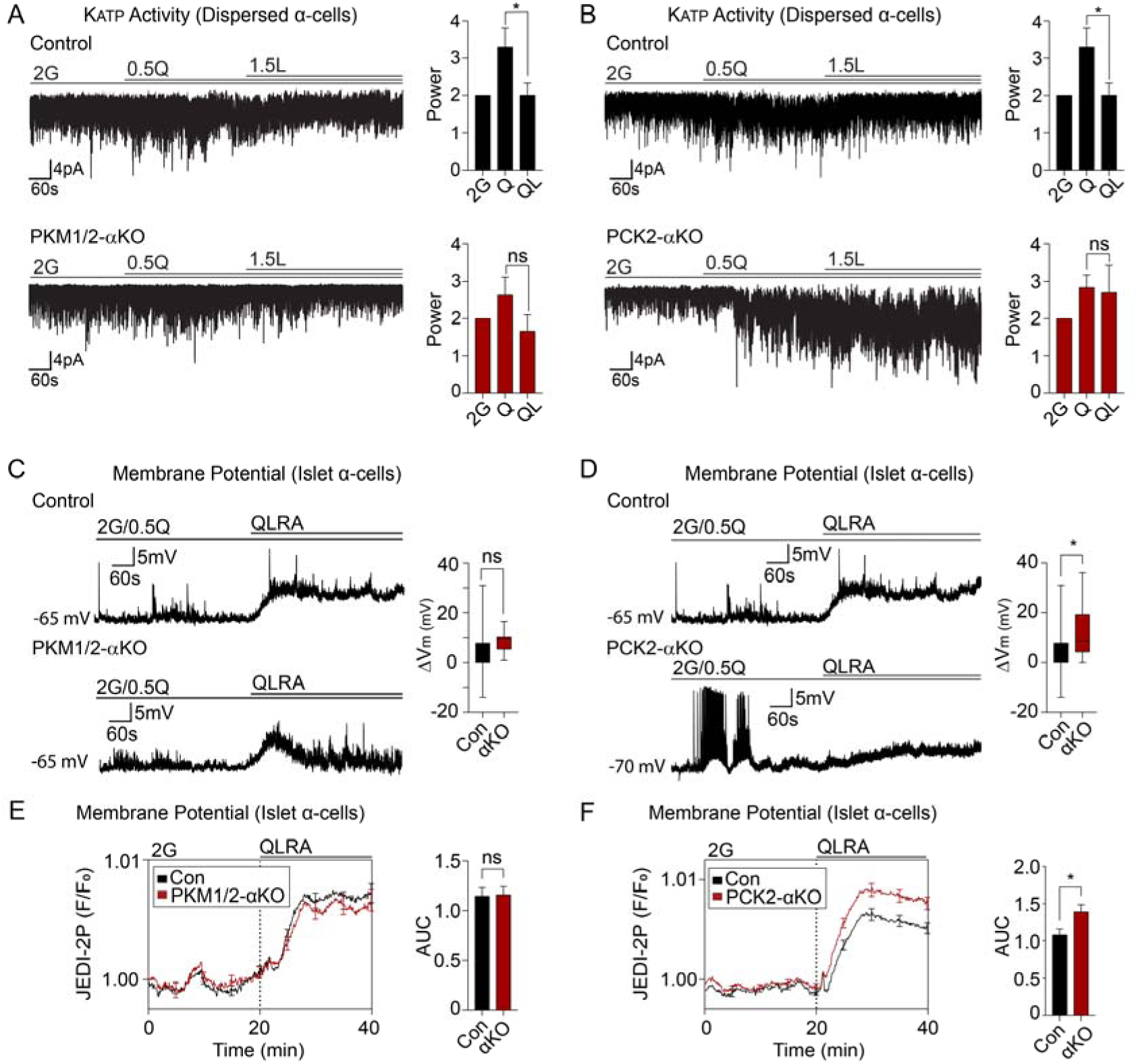
The α-cell PEP cycle closes K_ATP_ channels and restricts α-cell membrane potential in response to amino acid stimulation. *A-B*: Example traces and power analysis of K_ATP_ channel activity in response to 0.5 mM glutamine (0.5Q) and 1.5 mM leucine (1.5L) in dispersed α-cells from PKM1/2-αKO (A) and PCK2-αKO (B) and controls. *C-D*: Example traces and membrane potential (ΔV_m_) measured by patch-clamp in response 2 mM glucose and amino acids (QLRA; 0.6 mM glutamine, 0.5 mM leucine, 0.2 mM arginine, 2 mM and alanine) in intact islets from PKM1/2-αKO (*C*) and PCK2-αKO mice (*D*) as indicated. Data represent 12-18 cells from 3 mice per group. *E-F*: Average normalized membrane potential (ΔV_m_) measured by JEDI-2P voltage sensors in response to 2 mM glucose and QLRA in islets from PKM1/2-αKO (*E*) and PCK2-αKO mice (*F*) as indicated. Data represent 64-92 islets from 3 mice imaged per protocol. Data are shown as mean ± SEM for each condition compared with control by either by one way ANOVA (*A*, *B*) or Student’s *t* test (*C*-*F*). Controls were averaged in A-B and C-D. \**P* < 0.05, ns, not significant.

We next tested how PEP cycle affects α-cell V_m_ using perforated patch-clamp recordings of surface α-cells in the intact islet. α-Cells were identified by their characteristic I-V curves (Supplemental Fig. 2). The application of mixed amino acids (QLRA) depolarized α-cell V_m_ by 4.7 mV on average in control islets (Fig. 3C). Relative to controls, the amino acid-stimulated increase in ΔV_m_ trended upward in PKM1/2-αKO α-cells, but only reached significance in the PCK2-αKO (Fig. 3C,D). Given the high degree of α-cell variability, and the low throughput inherent to the patch-clamp approach, we performed orthogonal measurements using JEDI-2P voltage sensors (37) to record ΔV_m_ from hundreds of α-cells. In these assays, we isolated islets from PKM1/2-αKO and PCK2-αKO mice without GCaMP6s. Voltage-sensitive JEDI-2P biosensors (37) were expressed in α-cells using Cre-dependent adenovirus. Islets from PKM1/2-αKO and PCK2-αKO mice were measured simultaneously with their controls using barcoding with near-IR dyes. Confirming the patch-clamp studies, the α-cell ΔV_m_ response to amino acids was augmented by deletion of PCK2, while α-cell deletion of PKM1/2 had no significant effect (Fig. 3E,F).

### The PEP cycle suppresses **α**-cell Ca^2+^ in response to amino acid stimulation without impacting glucagon release

Our electrophysiological analyses of K_ATP_ channels and V_m_ suggested a mechanism by which PKM1/2 and PCK2 might regulate α-cell Ca^2+^. Consistently, the application of small molecule PK activators (10 μM TEPP-46), which allosterically activate PKM2 and PKL to drive the PEP cycle (34,36,38), nearly abolished the Ca^2+^ response to an amino acid ramp (Fig. 4A). To confirm that this effect was due to a direct effect on the α-cell, we applied the same amino acid ramp to islets isolated from PKM1/2-αKO and PCK2-αKO mice. Control islets were again measured simultaneously using islet barcoding. Elevated α-cell Ca^2+^ was observed in response to a range of amino acid levels in both PKM1/2-αKO and PCK2-αKO islets relative to controls (Fig. 4B,C). As in the electrophysiological assays, α-cell deletion of PCK2 had a stronger effect on Ca^2+^ levels than deletion of PKM1/2. Taking advantage of this phenotype, we used PCK2-αKO islets to test effects of individual amino acids on Ca^2+^ levels (Supplemental Fig. 3). Alanine and arginine were able to stimulate Ca^2+^ influx regardless of whether glutamine and leucine are present, as expected for their direct role in controlling V_m_. However, the Ca^2+^ phenotype of the PCK2-αKO islets was lost in the absence of glutamine, consistent with its role in providing anaplerotic glutamate carbons that drive oxaloacetate formation for PCK2. These data indicate that the α-cell PEP cycle cannot function in experiments lacking glutamine.

**Figure 4.**
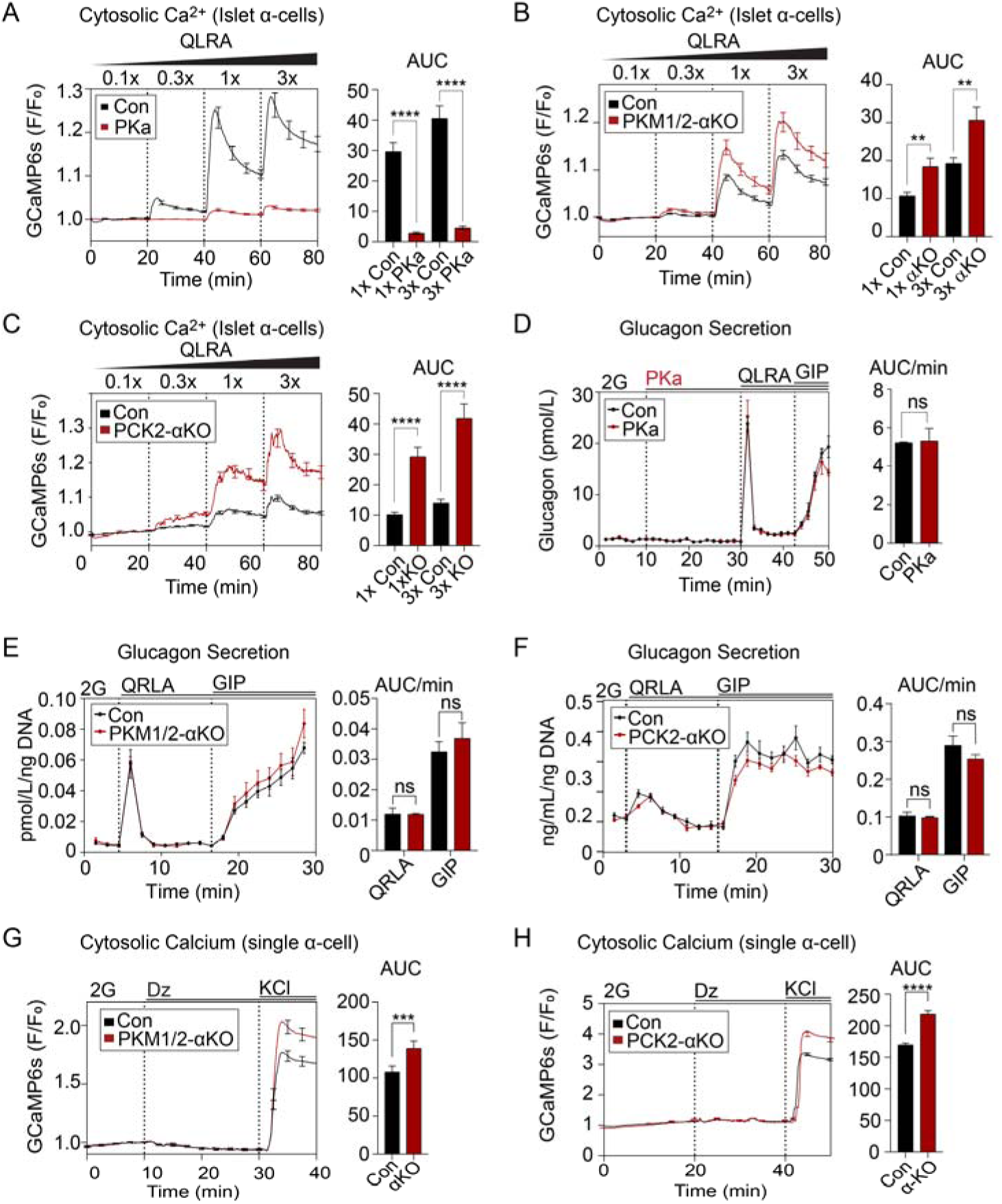
Pyruvate kinase and PCK2 suppress α-cell Ca^2+^ without impacting glucagon secretion. *A-C*: Average α-cell Ca^2+^ in response to amino acid ramp (1x QLRA; 0.6 mM glutamine, 0.5 mM leucine, 0.2 mM arginine, 2 mM and alanine) in islets from α-cell GCaMP6s mice treated with pyruvate kinase activator (PKa, 10µM TEPP-46), islets from control and PKM1/2-αKO mice (*B*), or control and PCK2-αKO mice (*C*). Data represent 57-105 islets from 3 mice per group. *D-E*: Average α-cell Ca^2+^ in response to 200 µM diazoxide followed by 30 mM KCl in intact islets from control and PKM1/2-αKO (*D*) and PCK2-αKO mice (*E*). Data represent 904-1194 α-cells from 4 mice. *F-H*: Glucagon secretion in response to QLRA, GIP (10 nM) in wild-type mice treated with PKa (*F*), control and PKM1/2-αKO mice (*G*), or control and PCK2-αKO mice (*H*). Data represent 6 chambers from 6 mice per condition with 80–100 mouse islets per chamber. Data are shown as mean ± SEM with AUC for each condition compared with control by Student’s *t* test. \*\**P* < 0.01, \*\*\**P* < 0.001, \*\*\*\**P* < 0.0001, ns, not significant.

We next investigated whether Ca^2+^ regulation by the PEP cycle impacts glucagon secretion using islet perfusion assays. Amino acids and GIP were sequentially applied under low glucose conditions to maximally stimulate glucagon release. However, we observed no difference in glucagon release upon PK activation with TEPP-46 (Fig. 4D), or upon α-cell deletion of PKM1/2 or PCK2 (Fig. 4E,F). Similar results were obtained in PCK2-αKO islets and controls were stimulated with alanine and GIP in the absence of leucine (Supplemental Fig. 4). The enhanced intracellular Ca^2+^ levels observed in α-cells lacking PKM1/2 and PCK2 is consistent with the permissive role of Ca^2+^ in controlling glucagon release. The ability of PK activation to suppress amino acid stimulated Ca^2+^ influx without impacting secretion not only indicates that Ca^2+^ influx at 2 mM glucose is sufficient to trigger glucagon secretion, but that amino acid stimulation of Ca^2+^ influx has no bearing on secretion.

Changes in cytosolic Ca^2+^ can also be modulated by Ca^2+^ extrusion. To test for a role of the PEP cycle downstream of V_m_ depolarization, we performed experiments in the presence of the K_ATP_ channel opener diazoxide (200 μM) and then triggered Ca^2+^ influx with 30 mM KCl. Even with V_m_ clamped, Ca^2+^ reached a higher level in α-cells lacking PKM1/2 or PCK2 (Fig. 4G,H). These findings indicate that the α-cell PEP cycle, in addition to controlling V_m_, homeostatically suppresses intracellular Ca^2+^ levels after glucagon is released.

## Discussion

Our data provide strong support for the concept of α-cells as amino acids sensors (3). We confirmed that alanine and arginine stimulate Ca^2+^ influx and glucagon secretion at physiological concentrations. Glutamine and leucine, while having only a small effect on Ca^2+^ at low glucose, strongly antagonized alanine/arginine-stimulated V_m_ depolarization and Ca^2+^ influx. We used three different models to show that glutamine and leucine suppress Ca^2+^ via α-cell PEP cycle, including pharmacological PK activators and α-cell specific deletion of PKM1/2 and PCK2. Strikingly, all three of these manipulations modulated Ca^2+^ without changing glucagon secretion. Thus, the glucagon released in response to amino acids, including electrogenic alanine and arginine, is not dependent on their ability to stimulate Ca^2+^ influx.

At the mechanistic level, we found that both PCK2 and PK are required for glutamine/leucine to close K_ATP_ channels, mirroring findings in β-cells (18,34–36), but the functional outcomes are different. While β-cells use the PEP cycle to initiate secretion, α-cells use it primarily to modulate Ca^2+^ dynamics. Specifically, we provided genetic evidence that PK is involved in suppressing intracellular Ca^2+^ levels in response to mitochondrial fuels. Part of this effect may arise from the ability of PK to close K_ATP_ channels and suppress membrane depolarization. These findings are consistent with our prior report indicating that a glycolytic metabolon is present on the plasma membrane of islet cells, and regulates α-cell K_ATP_ channels via the ATP/ADP ratio (18). Similar to glycolysis, mitochondrial PCK2 can generate PEP in response to glutamine/leucine. Deletion of PCK2 or PKM1/2 in α-cells blocked K_ATP_ closure in response to glutamine/leucine, indicating a conserved pathway in both α and β-cells (35). Of course, K_ATP_ closure has different effects on electrical excitability in α– and β-cells (11). In α-cells, we observed that both membrane potential and Ca^2+^ levels increased in response to amino acid stimulation in the PCK2 and PKM1/2 deletion models. In addition to K_ATP_ channels, there are other potential mechanisms of V_m_ regulation that we did not explore, such as metabolic control of α-cell Na^+^/K^+^ ATPases (39), which facilitate V_m_ hyperpolarization in β-cells (40).

Several factors may explain why elevated Ca^2+^ in our knockout models failed to enhance secretion. First is that Ca^2+^ is only permissive for hormone secretion in islet cells (6,41). From this perspective, it makes sense that elevated Ca^2+^ in the PCK2-αKO and PKM1/2-αKO has little impact on glucagon secretion, particularly in the presence of arginine and alanine that strongly stimulate Ca^2+^. Previous findings show that glucagon release is largely controlled by mouse α-cell P/Q-type Ca^2+^ channels, although they only account for ∼20% of Ca^2+^ current (10). Second, since glucagon release is not changed by PK or the PEP cycle, it is possible that K_ATP_ channels primarily impact L-type VDCC, as in β-cells. In any case, the field must look beyond Ca^2+^ to understand amino acid regulation of glucagon secretion. The glucagonostatic effect of leucine is both Ca^2+^– and PEP cycle-independent, and may be primarily mediated by its ability to reduce cAMP (31). Of note, gain-of-function mutations in glutamate dehydrogenase cause hyperinsulinemic hypoglycemia in response to protein rich meals (42). We believe this effect is due to glutamine carbons funneling through PCK2 and PK to close K_ATP_ channels (43). However, in α-cells, a change in glucagon secretion is not observed even when flux through PEP cycle is increased, potentially explaining why glutamate dehydrogenase mutations do not impact glucagon secretion.

Importantly, the elevated levels of Ca^2+^ in the knockout models were also observed in the presence of diazoxide/KCl, which stimulates Ca^2+^ influx independently of K_ATP_ closure. These findings indicate that the PEP cycle restores intracellular Ca^2+^ levels after depolarization. In cell lines, PK has been previously shown to play a role in ER Ca^2+^ storage (44). At low glucose, and in the absence of fatty acids (17), glutamine/leucine are key mitochondrial substrates. Defects in metabolism induced by α-cell PKM1/2 and PCK2 deletion could therefore impair Ca^2+^ extrusion. Mitochondria are also important for Ca^2+^ sequestration (45). In β-cells, the PEP cycle has been shown to hyperpolarize the mitochondrial membrane as PK lowers ADP (i.e., ADP privation) (34), and could affect mitochondrial Ca^2+^ handling. Further studies are needed to understand how amino acids affect organellar Ca^2+^ handling. In the meantime, our work shows that α-cells use the PEP cycle not as a direct regulator of secretion, but rather to limit Ca^2+^ rise during amino acid stimulation.

## Article Information

## Acknowledgements.

The authors thank the University of Wisconsin Carbone Cancer Center Flow Lab for assistance with cell sorting. The authors also thank Matthew Flowers from Dawn Davis lab from University of Wisconsin-Madison for assistance with qPCR. Graphics were created using BioRender.com.

## Funding

The Merrins laboratory gratefully acknowledges support from the NIH/NIDDK (R01DK113103 and R01DK127637 to M.J.M.) and the United States Department of Veterans Affairs Biomedical Laboratory Research and Development Service (I01BX005113 to M.J.M.). The Hughes laboratory gratefully acknowledges support from the NIH/NIDDK (R01DK138974 and R01DK140365 to J.W.H.).

## Duality of Interest

No potential conflicts of interest relevant to this article were reported.

## Authors contributions

E.J., H.R.F., E.P., S.M.H., X.D, and J.W.H. designed and performed experimental work and statistical analysis. E.J. and M.J.M. wrote and edited the manuscript. M.J.M. supervised the work.

## Supplementary data

**Supplemental Fig. 1.**
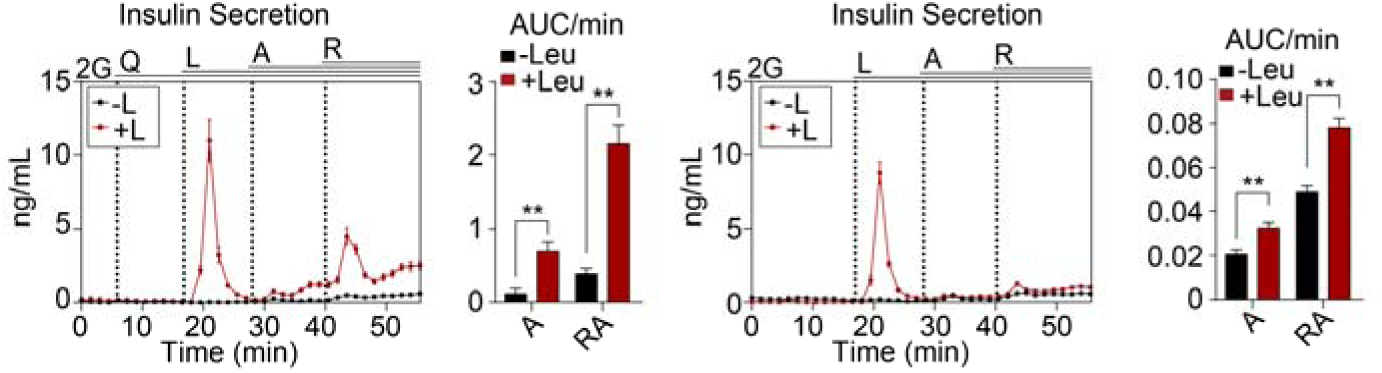
Leucine stimulates insulin secretion at low glucose. Insulin secretion from mouse islets measured as in Fig. 1I-J. Data are shown as mean ± SEM with AUC for each condition compared with control by Student’s *t* test. \*\**P* < 0.01.

**Supplemental Figure 2.**
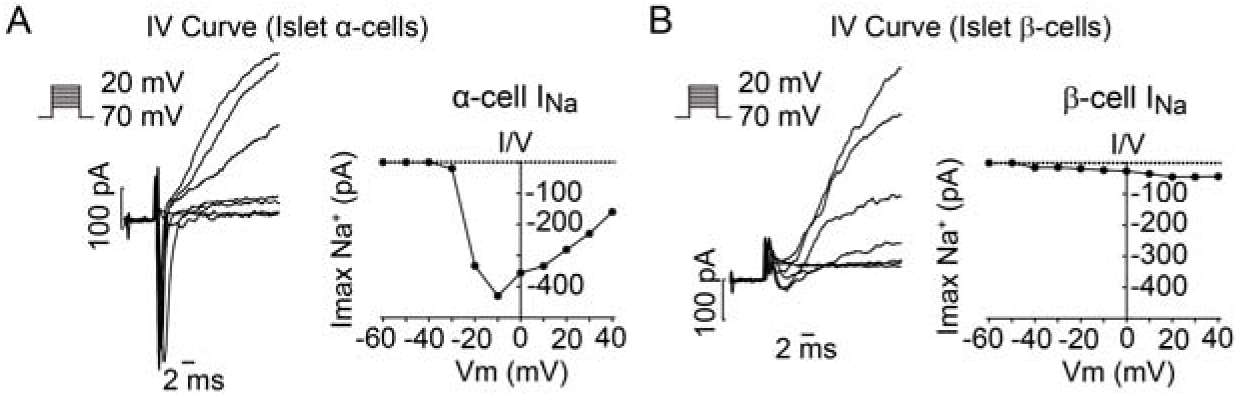
I–V curves used for α-cell identification in intact islets. Example traces and I-V curves measured from surface α-cells (*A*) and β-cells (*B*) in bath solution containing 2 mM glucose and 0.5 mM glutamine. Cells were depolarized from a holding potential of –70 mV to voltages between −60 and +40 mV in 10 mV increments. Data represent 12-18 cells from 3 mice per group as in Figure 3*C-D*.

**Supplemental Figure 3.**
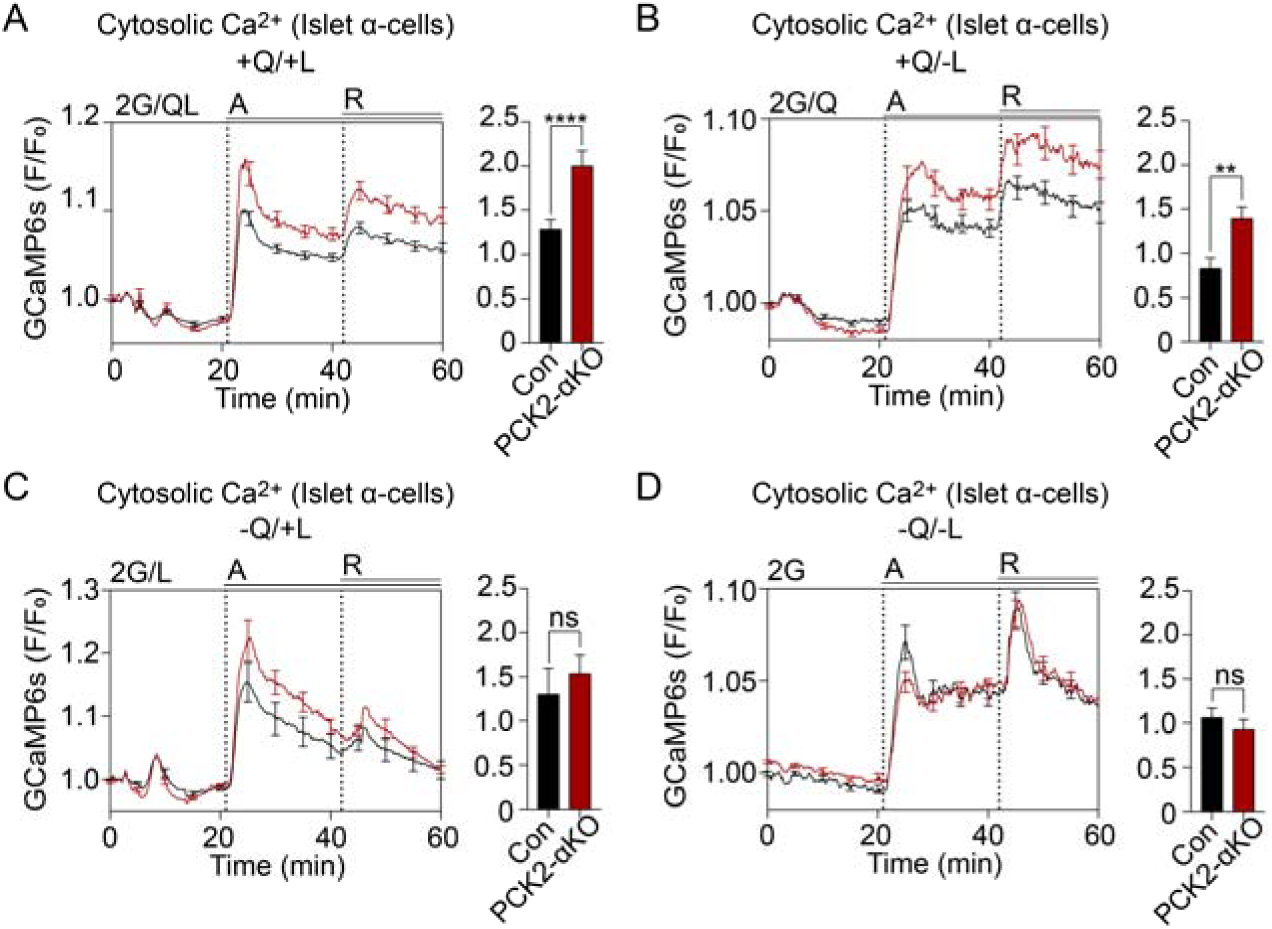
The α-cell Ca^2+^ PEP cycle requires glutamine. *A-D*: Averaged Ca^2+^ from control and PCK2-αKO mice in response to 2 mM glucose and combinations of amino acids as indicated. Data represent 42-55 islets from 3 mice per group. Data are shown as mean ± SEM with AUC for each condition compared with control by Student’s *t* test. \*\**P* < 0.01, \*\*\*\**P* < 0.0001, ns, not significant.

**Supplemental Figure 4.**
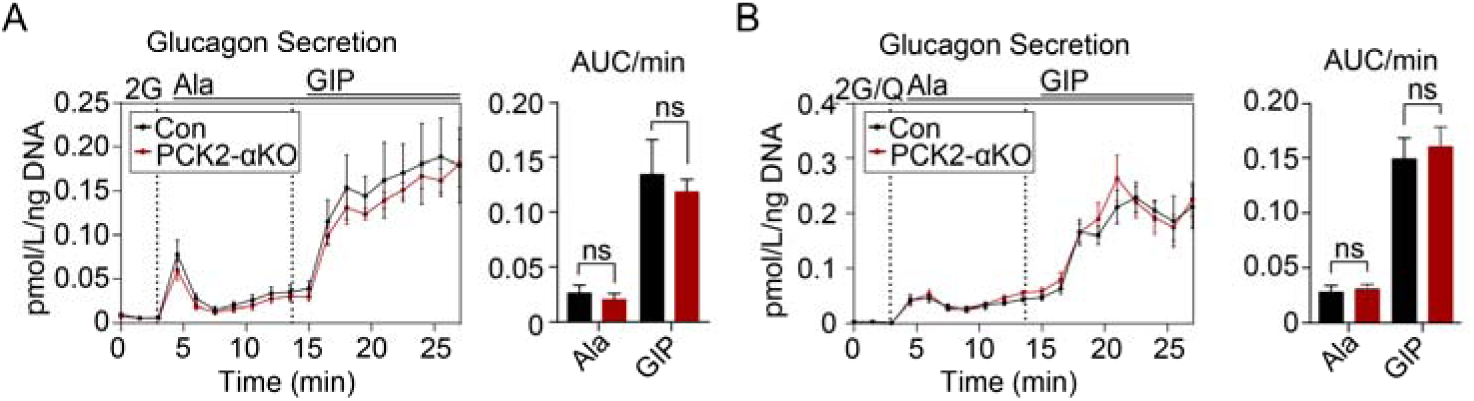
The α-cell Ca^2+^ PEP cycle is not required for glucagon secretion stimulated by alanine and GIP. Glucagon secretion control and PCK2-αKO islets in response to alanine (2 mM) and GIP (10 nM GIP). Data represent 3-6 chambers from 3-5 mice per condition with 80–100 mouse islets per chamber. Data are shown as mean ± SEM with AUC for each condition compared with control by Student’s *t* test. ns, not significant.

**Supplemental Figure 5.**
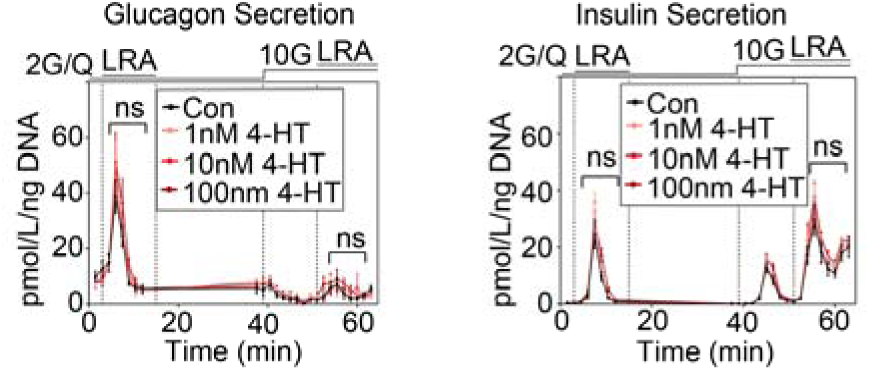
Culturing mouse islets 4-hydroxytamoxifen does not impact glucagon and insulin secretion. Hormone secretion measured from mouse islets after 3-day treatment with 1, 10, and 100 nM 4-hydroxytamoxifen (4-HT) in the culture media. Islets were measured in the presence of glucose (2G: 2mM glucose; 10G: 10mM glucose) and amino acids as indicated (QLRA, in mM: 0.6 mM glutamine, 0.5 mM leucine, 0.2 mM arginine, 2 mM and alanine). Data represent the average of 3 chambers from 3 mice per condition with 80 mouse islets per chamber. Data are shown as mean ± SEM with AUC for each condition compared with control by 2-way ANOVA. ns, not significant.

## Notes

### Competing Interest Statement

The authors have declared no competing interest.

### Summary of Updates

Added graphical abstract; Abstract and funding source edited

